# A deep CNN framework for neural drive estimation from HD-EMG across contraction intensities and joint angles

**DOI:** 10.1101/2022.01.17.476688

**Authors:** Yue Wen, Sangjoon J. Kim, Simon Avrillon, Jackson T. Levine, François Hug, José L. Pons

**Affiliations:** Legs and Walking Lab of Shirley Ryan AbilityLab and the Department of Physical Medicine and Rehabilitation, Feinberg School of Medicine, Northwestern University, Chicago, IL, USA; Nantes University, Laboratory ‘Movement, Interactions, Performance’ (EA 4334), Nantes, France; the University of Queensland, School of Biomedical Sciences, Brisbane, Australia; Institut Universitaire de France (IUF), Paris, France; and Université Côte d’Azur, LAMHESS, Nice, France; Legs and Walking Lab of Shirley Ryan AbilityLab, Department of Mechanical Engineering and Department of Biomedical Engineering, McCormick School of Engineering, and Department of Physical Medicine and Rehabilitation, Feinberg School of Medicine, Northwestern University, Chicago, IL, USA

**Keywords:** high-density electromyography (HD-EMG), neural drive, convolutional neural network (CNN), machine learning

## Abstract

**Objective:** Previous studies have demonstrated promising results in estimating the neural drive to muscles, the net output of all motoneurons that innervate the muscle, using high-density electromyography (HD-EMG) for the purpose of interfacing with assistive technologies. Despite the high estimation accuracy, current methods based on neural networks need to be trained with specific motor unit action potential (MUAP) shapes updated for each condition (i.e., varying muscle contraction intensities or joint angles). This preliminary step dramatically limits the potential generalization of these algorithms across tasks. We propose a novel approach to estimate the neural drive using a deep convolutional neural network (CNN), which can identify the cumulative spike train (CST) through general features of MUAPs from a pool of motor units.

**Methods:** We recorded HD-EMG signals from the gastrocnemius medialis muscle under three isometric contraction scenarios: 1) trapezoidal contraction tasks with different intensities, 2) contraction tasks with a trapezoidal or sinusoidal torque target, and 3) trapezoidal contraction tasks at different ankle angles. We applied a convolutive blind source separation (BSS) method to decompose HD-EMG signals to CST and segmented both signals into windows to train and validate the deep CNN. Then, we optimized the structure of the deep CNN and validated its generalizability across contraction tasks within each scenario.

**Results:** With the optimal configuration for the HD-EMG data window (overlap of 20 data points and window length of 40 data points), the deep CNN estimated the CST close to that from BSS, with a correlation coefficient higher than 0.96 and normalized root-mean-square-error lower than 7% with respect to the BSS (golden standard) within each scenario.

**Conclusion:** The proposed deep CNN framework can utilize data from different contraction tasks (e.g., different intensities), learn general features of MUAP variants, and estimate the neural drive for other contraction tasks.

**Significance:** With the proposed deep CNN, we could potentially build a neuraldrive-based human-machine interface that is generalizable to different contraction tasks without retraining.

## I. Introduction

The human-machine interface is a key component in wearable robots (i.e., prostheses and exoskeletons). Ideally, it should provide a seamless connection between the human and robot and achieve a reliable voluntary control of wearable robots in real time. When humans control their body, the motor cortex transmits neural signals to pools of motor units (MUs), including motor neurons and the muscle fibers they innervated, to generate force and movement [1]. Surface electromyography (EMG), a straightforward and easily accessed signal for intention detection, has been widely used for prostheses and exoskeleton control [2]–[5]. Surface EMG is the measurement at the surface of the skin of action potentials from muscle fibers activated by motor neurons [6]. However, the features (e.g., magnitude) of surface EMG signals cannot be used to accurately estimate the neural drive to muscles, the net output of all the motoneurons innervating the muscle as defined in [7], due to amplitude cancellation and cross-talk [6]. To address these limitations, recent studies relied on EMG decomposition methods to allow the extraction of spiking activities of MUs, which then can be used to calculate the cumulative spike trains (CST), an estimation of the neural drive [8]–[10]. Interestingly, it has been shown that using MU discharge timings to control upper-limb prosthesis outperformed conventional feature-based EMG control methods with both pattern recognition and musculoskeletal modelling [11].

There are two main approaches to use the neural drive to estimate muscle force or kinematics and generate motor commands. The first approach consists of estimating muscle force by summing the contribution of individual MUs based on optimization methods. Zhang et al. [12] and Yu et al. [13] have optimized a twitch model for each MU and summed the force outputs of all MUs to estimate muscle force. Alternatively, in a second approach, the smoothed CST can be used to estimate changes in isometric finger force [14], [15], finger kinematics [16], [17], and wrist kinematics [18], [19]. Importantly, the association of the smoothed CST with joint kinematics and kinetics using a simple linear regression can achieve performances comparable to the approaches based on individual MU spike trains, but with an easier implementation [14]–[19].

Current methods that estimate the neural drive are based on EMG decomposition, including gradient convolution kernel compensation (CKC) [8], FastICA peel-off [9], and convolutive blind source separation (BSS) [10]. The online version of these methods typically includes a two stage process: 1) an offline initialization where a separation matrix of MU filters is calculated based on a long segment of data (>10 s); 2) an online decomposition where the separation matrix is used to identify spike trains and then updated with the newly identified spike trains [14], [20], [21]. It is noteworthy that the number of MUs is determined by the separation matrix, which is fixed during the offline initialization. One limitation of these approaches is that the number of identifiable MUs may be different for different contraction intensities and even for similar intensities [22], [23]. In addition, the identifiable MUs change, especially when the contraction intensity increases [22], [24], due to amplitude cancellation of the smallest MUs [6]. Therefore, to the best of our knowledge, there is no approach that can be generalized across different contraction intensities to accurately estimate the neural drive.

Recently, researchers have demonstrated that a deep convolutional neural network (CNN) can estimate the wrist torque in two degrees of freedom with HD-EMG reconstructed using MU discharge times and the MUAP shapes. [13]. Deep recurrent neural networks (RNN) [25] and deep CNN [26] can also accurately identify the MU spike trains from HD-EMG. Specifically, these neural networks extract spatio-temporal features from the HD-EMG signal and then identify the spike train for individual MUs after a training based on EMG decomposition algorithms [25], [26]. However, these methods rely heavily on the features of the MUAP shapes, which are different for different MUs and could be altered by the muscle length even for the same MU [27].

Based on these promising results and the flexible structure of these neural networks, we propose a novel framework to directly identify the CST from HD-EMG without using individual MU filters. Thus, this would enable us to 1) combine training data from different contractions, either with identical or different intensities, without changing the structure of the neural network as we directly estimate the CST instead of a given number of MUs, 2) train the deep CNN to learn the features of the MUAP from all the identifiable MUs, and 3) utilize the deep CNN to identify the cumulative spike train in real time. In this study, we first investigate how the parameter space of the input (i.e., step size and window size) and output (i.e., number of output nodes) affect the performance of the deep CNN. Then, using the optimal parameters, we validate the generalizability of the deep CNN to estimate the neural drive of the gastrocnemius medialis (GM) under three scenarios: 1) isometric trapezoidal contractions with different intensities, 2) isometric contractions with a trapezoidal or sinusoidal torque target, 3) isometric contractions at different muscle length.

## II. Method

### A. Deep CNN model design

We have previously proposed a deep CNN approach to identify individual MU spike trains from HD-EMG signals [26]. In this previous study, the structure of the deep CNN was customized for each contraction intensity based on the number of MUs identified during preliminary training. Here, we propose to directly estimate the rate of spiking activities of the pool of MU, which is proportional to the neural drive, from HD-EMG signals by adapting the same deep CNN structure [26]. To this end, we modified the output of the deep CNN structure to identify the number of spikes in a given period of time (i.e., a window of HD-EMG signals). Moreover, this allowed us to create a unified structure that was generalizable to contractions with different number of MUs, such as contractions with different intensities.

Inspired in our prior work, here we used a window of HD-EMG signals as the input to the deep CNN, where the width of the window was M (number of HD-EMG channels) and the length was W (e.g., 120 data points from each channel when the window size was equal to 120). A sliding window approach adopted from typical EMG analysis methods [14], [15], [28] was used to segment the time-series HD-EMG signals into small windows of HD-EMG signals, and the increment of the sliding window was defined as step size (Fig. 1a). The output of the deep CNN was set to the number of MU spikes in a given period of time, as determined by the step size. If no spike was identified in the input window, the output of the deep CNN was zero for all the output nodes; if one spike was identified in the window, the output was set to one in the first output node; if two spikes were identified, the output was set to one in the two first output nodes; and so on.

**Fig. 1:**
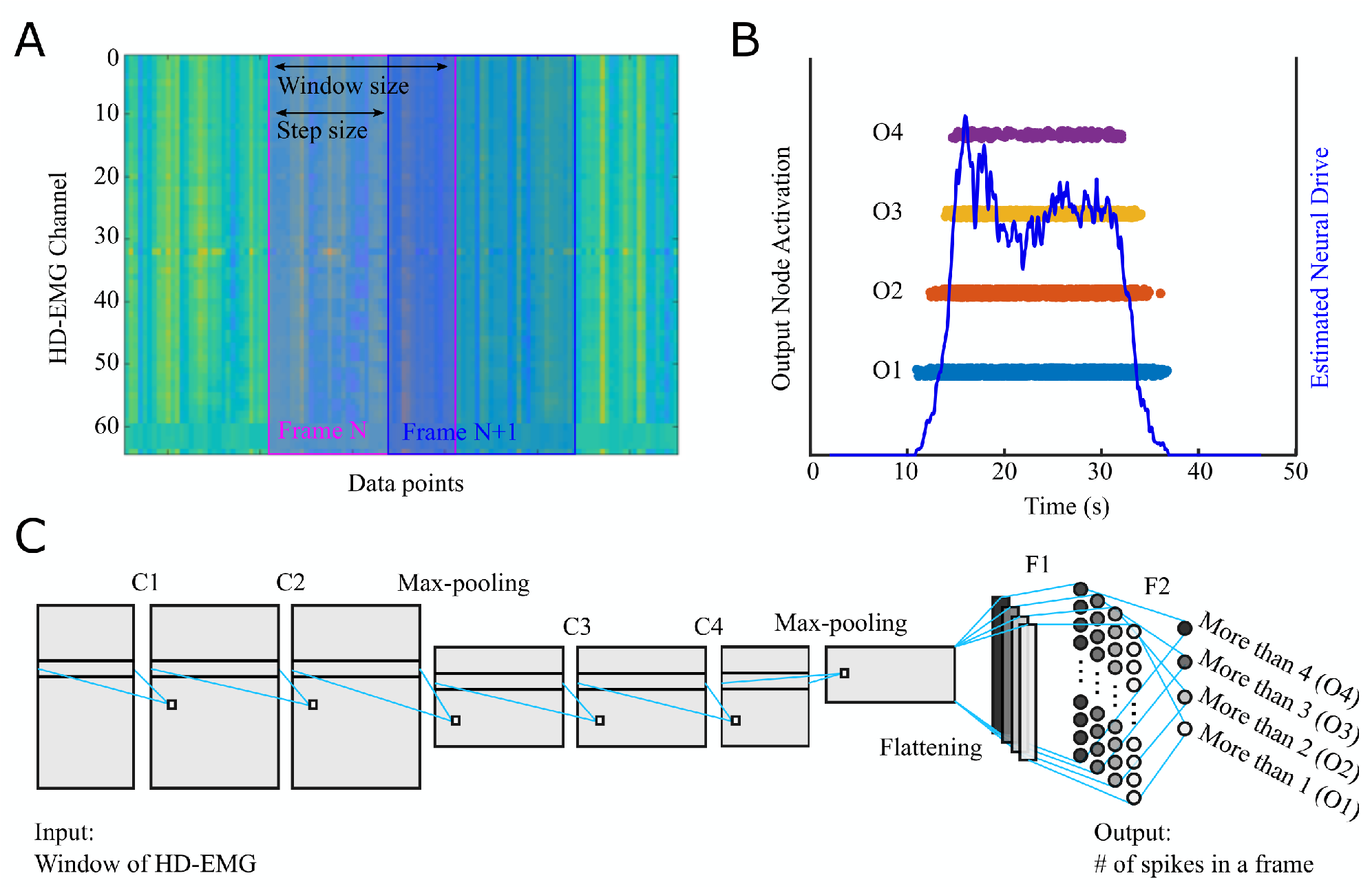
Overview of the deep CNN framework. Panel (A) represents two consecutive windows of HD-EMG signals as inputs to the deep CNN, (B) the activation of the output nodes and the estimated neural drive in form of smoothed CST, and (C) the overall structure of the deep CNN.

We implemented six parametric layers (four convolutional layers and two fully connected layers) and three non-parametric layers (two max-pooling layers and one flatten layer) between the input layer and output layer (Fig. 1c). The first three convolutional layers (C1–C3) included 128 parallel kernel filters with a kernel size of 3 to extract the spatio-temporal features, while the last convolutional layer (C4) had 64 kernel filters to decrease the number of trainable parameters of the fully connected layers. The output of the convolutional layers was flattened to a vector and fed into two fully connected layers, including 256 nodes and 1 node, respectively. We used a sigmoid activation function for the output layer and a rectified linear unit (ReLU) activation function for all other layers. The sigmoid activation function had an output ranging from 0 to 1, which adhered to our output requirement of being a binary value. The ReLU helped to solve the vanishing gradient problem and accelerate the convergence speed. We have designed the consecutive convolutional layers (C1 and C2, C3 and C4) to allow the CNN model to learn more complex features. Max-pooling with size of 2 was applied to C2 and C4 to extract representative features while reducing the dimension of the features. Then, the fully connected layers made predictions based on the feature maps from convolutional layers. In addition, a dropout rate of 50% was applied to C2, C4, and F2 layers to regularize the neural network and to prevent over-fitting.

### B. Experimental signals

Three experimental datasets were recorded from the GM muscle and used in this study: 1) isometric trapezoidal contractions with three different intensities (Fig. 3A), 2) isometric contractions tracking a trapezoidal or sinusoidal torque target (Fig. 3B), and 3) isometric trapezoidal contractions at different ankle angles (Fig. 3C). Such experimental design is to consider the variants of the MUAP shapes and MU firing patterns when different MUs are recruited at different contraction intensities/targets or when muscle length is changed at different joint angles. The first dataset was used to optimize the input and output parameters of the deep CNN. Then, all three datasets were used to investigate the generalization of the proposed deep CNN approach within these three scenarios.

**Fig. 2:**
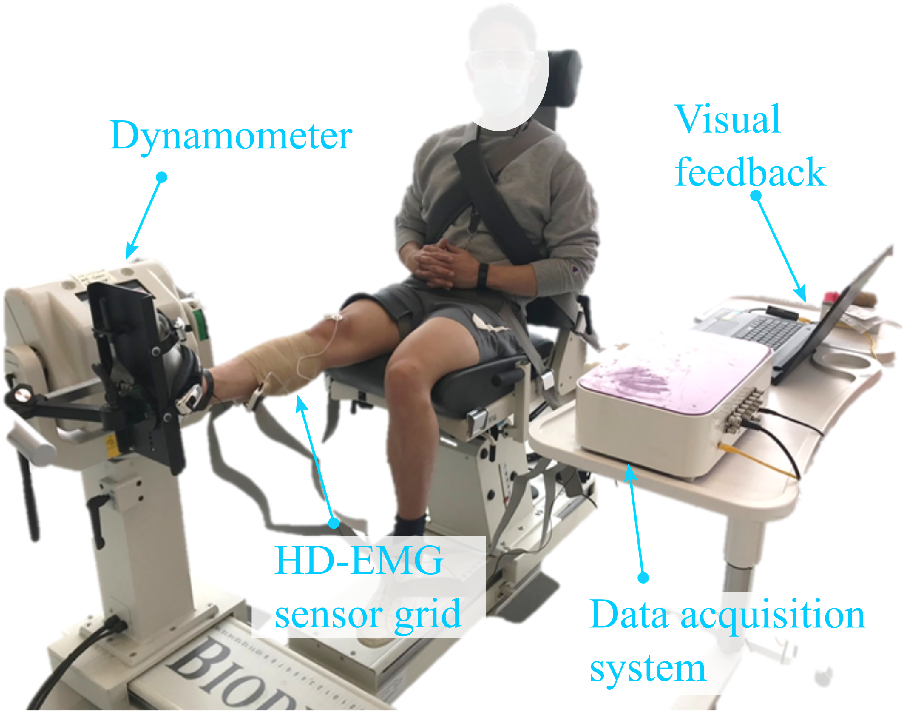
Experimental setup for the generalization experimentation across torque targets and ankle angles (experiment 2 and 3).

**Fig. 3:**
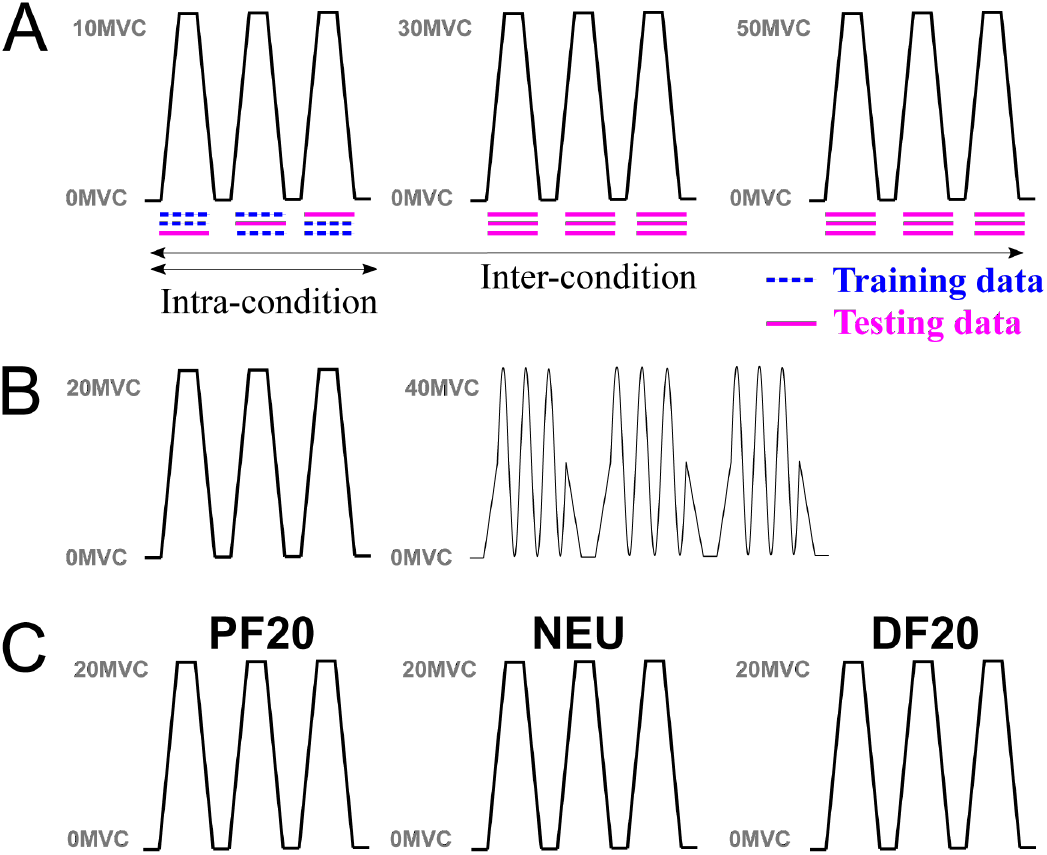
Experimental protocol to train and validate the deep CNN. Panel (A) is for isometric trapezoidal contractions with different intensities: 10% MVC, 30% MVC, and 50% MVC, (B) isometric contractions with different torque targets: trapezoidal and sinusoidal targets, and (C) isometric trapezoidal contractions at different ankle angles: plantarflexion at 20 degrees (PF20), neutral posture (NEU), and dorsiflexion at 20 degrees (DF20). An example of the training and validation protocol is presented in panel (A). The blue dotted lines represent the data used to train the deep CNN and the solid pink lines represent the data used to validate the deep CNN.

For all experiments, a two-dimensional adhesive grid of 64 electrodes (13 × 5 electrodes with one electrode absent on a corner, gold-coated, inter-electrode distance: 8 mm; [ELSCH064NM2, SpesMedica, Battipaglia, Italy] for the first experiment, and [GR08MM1305, OT Bioelettronica, Italy] for the other experiments) was placed over the GM muscle. Before applying the electrode, the skin was shaved and cleaned with an abrasive pad and alcohol. The adhesive grid was attached on to the skin using semi-disposable bi-adhesive foam layers (SpesMedica, Battipaglia, Italy). Skin to electrode contact was made by filling cavities of the adhesive layers with conductive paste (SpesMedica, Battipaglia, Italy). After attaching the electrodes to the skin, an elastic band was placed over the electrode to ensure good contact throughout the experiment. A strap electrode dampened with water was placed around the contralateral ankle as a reference electrode. For experiment 1, another dampened strap electrode was placed around the ipsilateral ankle as a ground electrode; for experiments 2 and 3, a sticky ground electrode was placed on the bony area of the ipsilateral tibia. The EMG signals were recorded in monopolar mode, band-pass filtered (10–900 Hz), and digitized at a sampling rate of 2048 Hz using a multichannel acquisition system (EMG-Quattrocento; 400-channel EMG amplifier, OT Bioelettronica, Italy). The torque signal was low-pass filtered (Butterworth 3rd order, cut-off frequency: 20 Hz) and digitized at 2048 Hz using the same acquisition system that was used for HD-EMG.

#### 1) Isometric trapezoidal contractions with different intensities

Six healthy participants have participated in this study (6 males, 34±8 years, height 178±6 cm, body mass 76±7 kg). All participants had no history of lower leg injury within the past 6 months. The institutional research ethics committee of the University of Queensland approved this study (No. 2013001448), and all procedures were in accordance with the Declaration of Helsinki. All participants provided their written informed consent prior to participation in the study. Note that the experimental data were a subset of a previously published dataset [29].

After placing the electrodes, participants were asked to lay prone on a custom dynamometer while positioning their knee in full extension. The custom dynamometer, which consisted of a foot pedal and a torque sensor (TRE-50K, Dacell, Rep. of Korea) was used to collect data. The ankle of the participant was securely fastened at a plantarflexion angle of 10 degrees (0 degree being the foot perpendicular to the shank). After performing a warm-up set of a series of submaximal contractions, participants performed three maximal isometric voluntary contractions (MVC) for 3–5 s with 120 s rest between contractions to prevent muscle fatigue. Then, participants performed three contractions at each of the following intensities: 10%, 30%, and 50% of their MVC. The order of the intensities was randomized. These contractions involved a 5 s ramp-up, 20 s (10% and 30% MVC) or a 15 s (50% MVC) plateau, and a 5 s ramp-down phase. The contractions were separated by either 60 s (10% MVC) or 120 s (30 and 50% MVC) to reduce the effect of muscle fatigue. Both HD-EMG and torque signals were recorded concurrently using aforementioned acquisition system.

#### 2) Isometric contractions with trapezoidal or sinusoidal torque targets

Three healthy participants have participated in this study (3 males, 27±3 years, height 181±6 cm, body mass 71±13 kg). All participants had no history of lower leg injury within the previous 6 months that could prevent their full engagement during the experiments. The institutional research ethics committee of Northwestern University approved this study (No. STU00212191), and all procedures were in accordance with the Declaration of Helsinki.

After placing the electrodes, participants were asked to sit upright on a dynamometer (Biodex, USA) while positioning their knee in full extension (see Fig.2). The trunk, thigh and ankle of the participant were securely fastened to the dynamometer. The ankle was fixed at a neutral position where the foot was perpendicular to the shank. After performing a warm-up set of a series of sub-maximal contractions, participants performed two isometric MVC for 3–5 s with 120 s rest in between.

After the MVC contractions, participants performed isometric contractions during which they tracked different torque targets: a trapezoidal target and a sinusoidal target. As depicted in Fig. 3B, the trapezoidal contraction involved a 15 s rampup, a 30 s plateau at 20% MVC, and a 15 s ramp-down phase, while the sinusoidal contraction involved a 15 s ramp-up, 30 s of sine wave tracking in the range of 0-40% MVC with a frequency of 0.1 Hz, and a 15 s ramp-down phase. For each torque target, the participants performed three contractions, and, in between contractions, they took 60-s rest to avoid the occurrence of muscle fatigue. During the experimentation, both HD-EMG and torque signals were recorded concurrently using aforementioned acquisition system.

#### 3) Isometric trapezoidal contractions with different ankle angles

Three healthy participants participated in this experiment (3 males, 29±6 years, height 183±3 cm, body mass 79±9 kg). They met the same criteria and signed the same consent form (approved by research ethics committee of the Northwestern University) as mentioned in experiment 2. All preparation procedures and the trapezoidal torque target were the same as experiment 2.

After preparation and MVC contractions as in experiment 1, participants performed three isometric contractions at 20% MVC with each being performed at a different ankle angle (Fig. 3C): 1) 20 degrees plantarflexion (PF20), 2) a neutral posture (NEU), and 3) 20 degrees dorsiflexion (DF20). During the experimentation, both HD-EMG and torque signals were recorded concurrently using aforementioned acquisition system.

### C. Data processing

#### 1) HD-EMG decomposition with the BSS algorithm

The EMG signals from all channels were first band-pass filtered (Butterworth 2nd order, 20–500 Hz). Then they were visually inspected and the noisy channels with low signal-to-noise ratio or motion artifacts were removed. After cleaning the data, the HD-EMG signals were decomposed into motor unit spike trains using a convolution blind-source separation method [10], which has been validated using experimental and simulated signals [10]. After the automatic identification of the MUs, all the MU spike trains were visually inspected and manually edited when a false positive (FP; labeled artifact) or a false negative (FN; non-labeled discharges) was observed by an experienced operator. Only the MUs that exhibited a pulse-to-noise ratio (PNR) >30 dB were retained for further analysis. This threshold ensured a sensitivity higher than 90% and a FP rate lower than 2% [30]. In addition, MU spike trains with less than 150 spikes were not considered for the analysis [26].

#### 2) Deep CNN parameter optimization

As mentioned in Section II.A, we fixed the internal structure of the deep CNN but allowed flexible input and output structure for this HD-EMG based neural drive estimation problem. We optimized the input and output parameters with a dataset including isometric trapezoidal contractions with three different intensities (i.e., 10% MVC, 30% MVC, 50% MVC).

The optimization procedure included two steps: 1) we first investigated the effects of input parameters (i.e., step size and window size). Based on our previous study [26], we considered 16 combinations of 4 step size conditions (i.e, ST5, ST10, ST20, ST40) and 4 window size conditions (i.e., WS20, WS40, WS80, and WS120) and 3 nodes in the output layer; 2) with the top two performing combinations of the step size and window size, we further investigated the effects of the number of output nodes ranging from 1 to 6. This resulted in 12 combinations from 2 input conditions and 6 output conditions. First, the data (three contractions for three intensities) were reorganized to three folds of data, each fold including one contraction from each intensity. Then, we performed three-fold cross-validation [26], using two folds as training data and the remaining fold as testing data, to assess the performance of deep CNN for each combination of window size, step size, and number of output nodes. The performance was averaged across three validation runs for each combination. Lastly, we calculated the mean and standard deviation of the performance of the deep CNN across all participants for each combination of step size, window size, and number of output nodes.

#### 3) Generalizability and scalability of the deep CNN

The generalizability of the deep CNN was defined as the performance of the deep CNN when applied to data from tasks other than those of training data (details in Section II.D). As shown in Fig. 3, we validated the generalizability of the deep CNN within each one of the three scenarios: 1) across contractions with different intensities (i.e., 10% MVC, 30% MVC, and 50% MVC), 2) across contractions with different targets (i.e., trapezoidal and sinusoidal torque targets), 3) across contractions at different ankle angles (i.e., PF20, NEU, and DF20). Within each scenario, we performed intra-task validation, where the training and testing data were from the same contraction task, and inter-task validation, where the training and testing data were from different contraction tasks. Furthermore, we combined datasets from different contraction tasks to validate the scalability of the deep CNN to such complex situations.

Taking scenario 1 as an example (Fig. 3A), three repeated contractions for each intensity generated three groups/folds of data. To evaluate the deep CNN when trained with data from trapezoidal contraction with 10% MVC, a three-fold cross-validation was performed. For intra-task validation, two folds of the data (blue dash lines in Fig. 3A) were used to train the deep CNN while the remaining one (pink line in Fig. 3A) was used to test the performance of the deep CNN (left panel of Fig. 3A). This generated three trained deep CNNs. For intertask validation, the three trained deep CNNs were applied to all three folds of data from a different contraction intensity (i.e., 30% MVC or 50% MVC; middle and right panels of Fig. 3A). Lastly, the performance of the deep CNN was averaged across three tests for intra-task validation and across nine tests for inter-task validation (pink lines in Fig. 3A). The same intra-task and inter-task validations were performed for each contraction task in each scenario.

In addition, we combined the data from all contraction tasks in each scenario to form three folds of data, each fold including one contraction from each contraction task (e.g., one from each contraction intensity), and then performed another three-fold cross-validation. The purpose was to assess the scalability of the deep CNN to a more complex situation with combined contraction tasks.

### D. Evaluation criteria

The outputs of the deep CNN were summed to form the CST, which was then smoothed using a 400-ms Hanning window to estimate the neural drive [31].

We calculated the correlation coefficient and the normalized root-mean-square error (nRMSE) between the estimated neural drive from manually edited spike trains from BSS and the neural drive estimated from the deep CNN to evaluate the accuracy in neural drive estimation. Specifically, the nRMSE was defined as

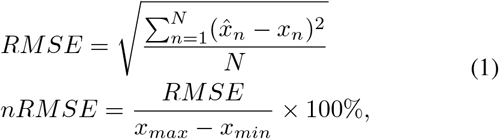

where 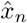 and *x_n_* are the nth sample of the smoothed neural drive from deep CNN and the smoothed neural drive from BSS, respectively. *x_max_* and *x_min_* are the maximum and minimum values of the smoothed neural drive from BSS, and N is the number of samples.

As the deep CNN could generate false positives while the muscle was not activated, we included 2-s of data before and after the actual contraction when calculating the correlation coefficient and nRMSE.

## III. Results

### A. Motor unit decomposition using BSS

With the BSS method and manual editing, we identified, across all contraction tasks and subjects in each scenario, 20±6 MUs (range 8-30) with 12269±3715 spikes in scenario 1, 29±5 MUs (range 23-35) with 16616±5462 in scenario 2, and 19±7 MUs (range 7-29) with 31754±9261 spikes in scenario 3. Table. I shows the detailed numbers of MUs and spikes per participant for each contraction task.

**TABLE I:** The number of MUs and spikes decomposed using BSS for each contraction task across all participants.

|  | # of MUs<sup>a</sup> | # of spikes<sup>b</sup> |
| --- | --- | --- |
| 10% MVC | $22 \pm 5$ | $11794 \pm 2738$ |
| 30% MVC | $23 \pm 4$ | $16293 \pm 3208$ |
| 50% MVC | $14 \pm 5$ | $10087 \pm 3221$ |
| TRAP | $22 \pm 6$ | $20681 \pm 1082$ |
| SINE | $17 \pm 9$ | $12550 \pm 4881$ |
| DF20 | $28 \pm 6$ | $29188 \pm 6022$ |
| NEU | $29 \pm 6$ | $31327 \pm 10135$ |
| PF20 | $29 \pm 6$ | $34748 \pm 13434$ |

<sup>a</sup> the number of MUs per participant. <sup>b</sup> the number of spikes per participant.

### B. Optimization of the network input and output structure

Among the 16 combinations from 4 step sizes (i.e., ST5, ST10, ST20, and ST40) and 4 window sizes (i.e., WS20, WS40, WS80, and WS120), the deep CNN generated correlation coefficients higher than 0.97 and nRMSE lower than 6.6% for ST10-WS120, ST20-WS40, ST20-WS80, and ST20-WS120 (Table II). The highest correlation coefficient (0.97±0.01) and lowest nRMSE (6.3±1.0%) were achieved for ST20-WS40. The effect of number of outputs was then validated with the two highest performing window size and step size combinations (i.e., ST20-WS40 and ST10-WS120). For both combinations, when the number of outputs increased from one to six, the correlation coefficient increased (Fig. 4A) and the nRMSE decreased (Fig. 4B), indicating the deep CNN generated increasingly more accurate estimation of neural drive. The correlation coefficient and nRMSE saturated when the number of outputs was set to four for both ST20-WS40 and ST10-WS120. When comparing between ST10-WS120 and ST20-SW40, the highest performance with the four output configuration was achieved for ST20-WS40 (Fig. 4).

**TABLE II:** The correlation coefficient and nRMSE of the deep CNN with different window size and step size.

| r | WS20 | WS40 | WS80 | WS120 |
| --- | --- | --- | --- | --- |
| ST5 | $0.53 \pm 0.23$ | $0.85 \pm 0.14$ | $0.83 \pm 0.25$ | $0.82 \pm 0.23$ |
| ST10 | $0.86 \pm 0.06$ | $0.96 \pm 0.02$ | $0.96 \pm 0.02$ | $0.97 \pm 0.02$ |
| ST20 | $0.96 \pm 0.02$ | $0.97 \pm 0.01$ | $0.97 \pm 0.02$ | $0.97 \pm 0.01$ |
| ST40 | $0.93 \pm 0.03$ | $0.94 \pm 0.02$ | $0.94 \pm 0.02$ | $0.95 \pm 0.02$ |
| nRMSE | - | - | - | - |
| ST5 | $20.5 \pm 3.4$ | $11.8 \pm 3.7$ | $11.6 \pm 6.1$ | $12.3 \pm 6.2$ |
| ST10 | $12.4 \pm 2.6$ | $7.1 \pm 1.5$ | $6.6 \pm 1.6$ | $6.3 \pm 1.3$ |
| ST20 | $7.0 \pm 0.9$ | $6.3 \pm 1.0$ | $6.6 \pm 1.0$ | $6.5 \pm 0.7$ |
| ST40 | $9.3 \pm 1.2$ | $8.6 \pm 0.9$ | $8.2 \pm 1.1$ | $8.3 \pm 1.1$ |

The nRMSE is in %.

**Fig. 4:**
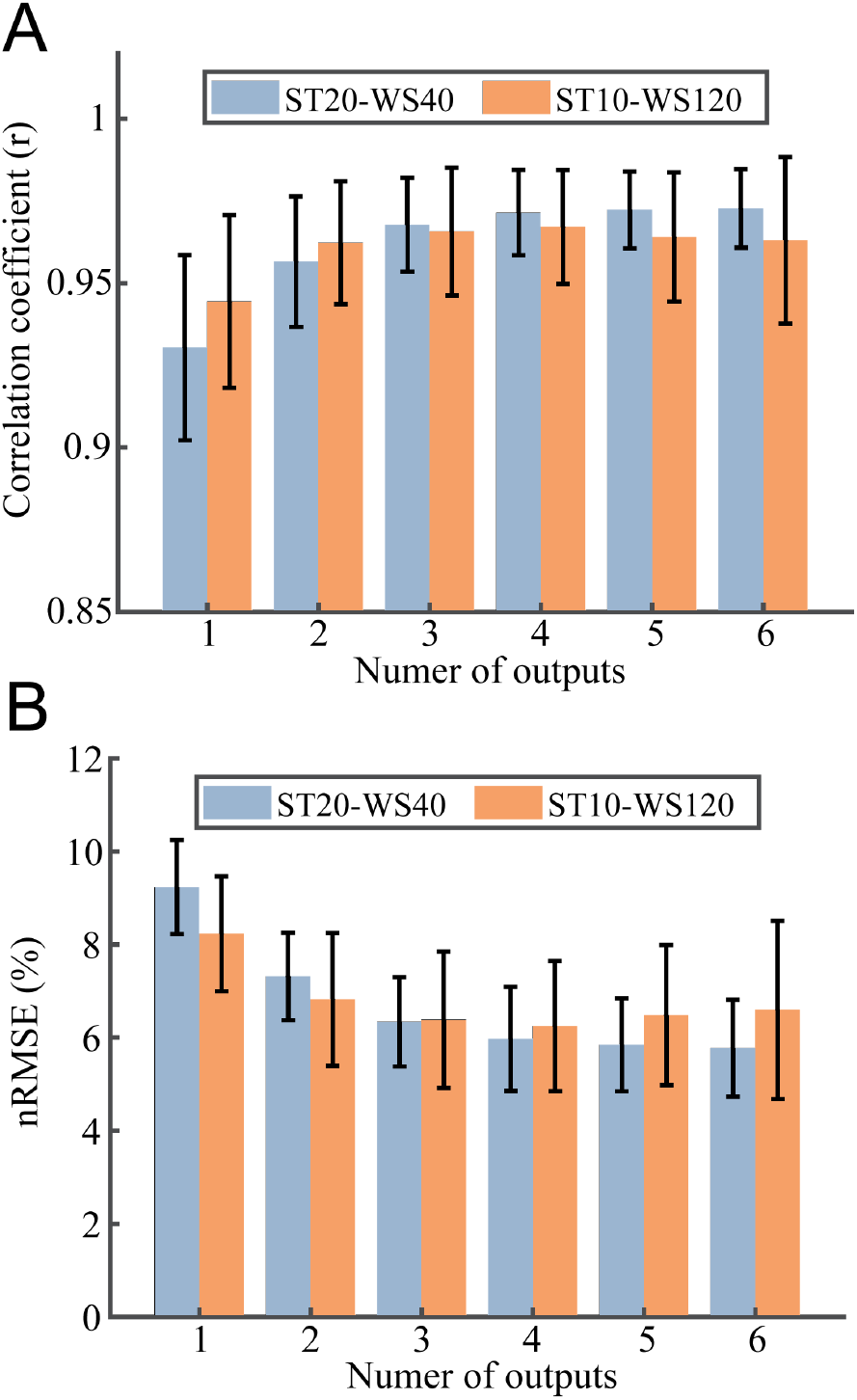
The correlation coefficient and nRMSE of the deep CNN with different number of outputs in combination with two input formats: ST20-WS40 and ST10-WS120. ST and WS denoted the step size and window size in windowing the HD-EMG signals, respectively. The error bar was calculated across six participants.

### C. Evaluation of the deep CNN for neural drive estimation

Table III, IV, and V present the correlation coefficient and nRMSE of the deep CNN for all validations across contractions with different intensities, with different torque targets,

**TABLE III:**
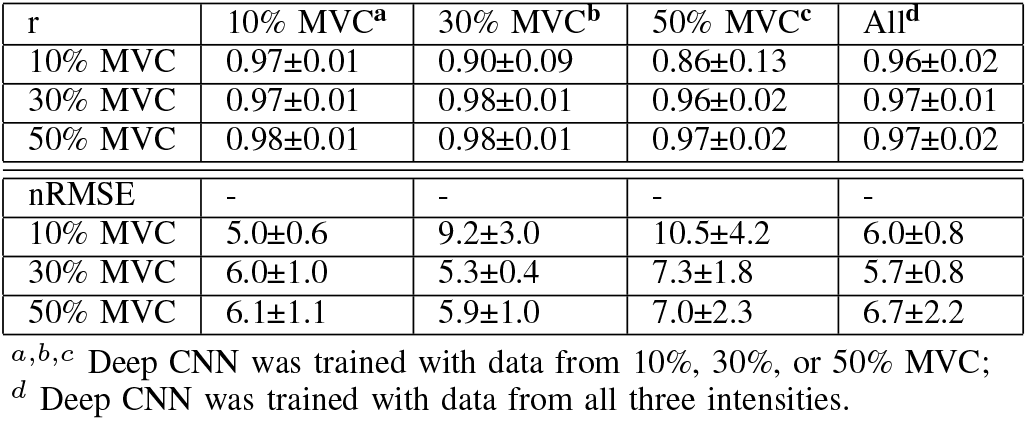
The correlation coefficient and nRMSE of deep CNN trained and validated across contractions with different intensities

**TABLE IV:** The correlation coefficient and nRMSE of the deep CNN trained and validated across contractions with trapezoidal and sinusoidal torque targets.

| r | 20% MVC<sup>a</sup> | Sine<sup>b</sup> | All<sup>c</sup> |
| --- | --- | --- | --- |
| Trap | 0.98±0.01 | 0.96±0.02 | 0.97±0.01 |
| Sine | 0.97±0.01 | 0.98±0.01 | 0.98±0.02 |
| nRMSE | - | - | - |
| Trap | 5.73±0.88 | 7.46±2.02 | 5.93±0.94 |
| Sine | 6.32±1.76 | 5.29±1.80 | 5.53±2.39 |

<sup>a,b</sup> Deep CNN was trained with data from trapezoidal torque tracking or sinusoidal torque tracking;<sup>c</sup> Deep CNN was trained with data from both contraction tasks.

**TABLE V:** The correlation coefficient and nRMSE of the deep CNN trained and validated across contractions at different ankle angles.

| r | PF20<sup>a</sup> | NEU<sup>b</sup> | DF20<sup>c</sup> | All<sup>d</sup> |
| --- | --- | --- | --- | --- |
| PF20 | 0.97±0.01 | 0.97±0.01 | 0.95±0.02 | 0.97±0.01 |
| NEU | 0.98±0.01 | 0.98±0.01 | 0.97±0.01 | 0.98±0.01 |
| DF20 | 0.96±0.01 | 0.96±0.01 | 0.98±0.01 | 0.98±0.01 |
| nRMSE | - | - | - | - |
| PF20 | 6.20±0.47 | 6.38±0.61 | 8.16±2.05 | 6.04±0.53 |
| NEU | 5.84±0.75 | 5.43±0.67 | 6.55±1.52 | 5.40±0.65 |
| DF20 | 6.93±0.37 | 6.66±0.80 | 5.25±0.24 | 5.26±0.58 |

and at different ankle angles, respectively. For across intensity validation (Table III), the correlation coefficient and nRMSE were 0.97±0.01 and 5.78±1.07 for intra-task validation (i.e., the diagonal elements with training and testing data from same contraction task), and they changed to 0.94±0.05 and 7.50±1.94, respectively, for inter-task validation (i.e., the non-diagonal elements in the first three columns with training and testing data from different contraction tasks). For across tracking target validation (Table IV), the correlation coefficient and nRMSE were 0.98±0.00 and 5.51±0.31 for intra-task validation, and they changed to 0.97±0.00 and 6.89±0.04 for inter-task validation. Fig. 5 illustrates a representative neural drive comparison between BSS (red line) and deep CNN (blue line) for intra-task and inter-task validations across trapezoidal and sinusoidal torque targets. For across joint angle validation (Table V), the correlation coefficient and nRMSE were 0.98±0.01 and 5.63±0.50 for intra-task validation, and they changed to 0.97±0.01 and 6.75±0.78 for inter-task validation. Fig. 6 illustrates the neural drive comparison for intra-task and inter-task validations with different ankle angles. For validation with combined data (i.e., training and testing with combined data from different contraction tasks), the correlation coefficient and nRMSE (last column in Table III, IV, and V) were 1) 0.97±0.01 and 6.13±0.51 for scenario 1, respectively; 2) 0.98±0.01 and 5.73±0.28 for scenario 2; and 3) 0.98±0.01 and 5.57±0.42 for scenario 3.

**Fig. 5:**
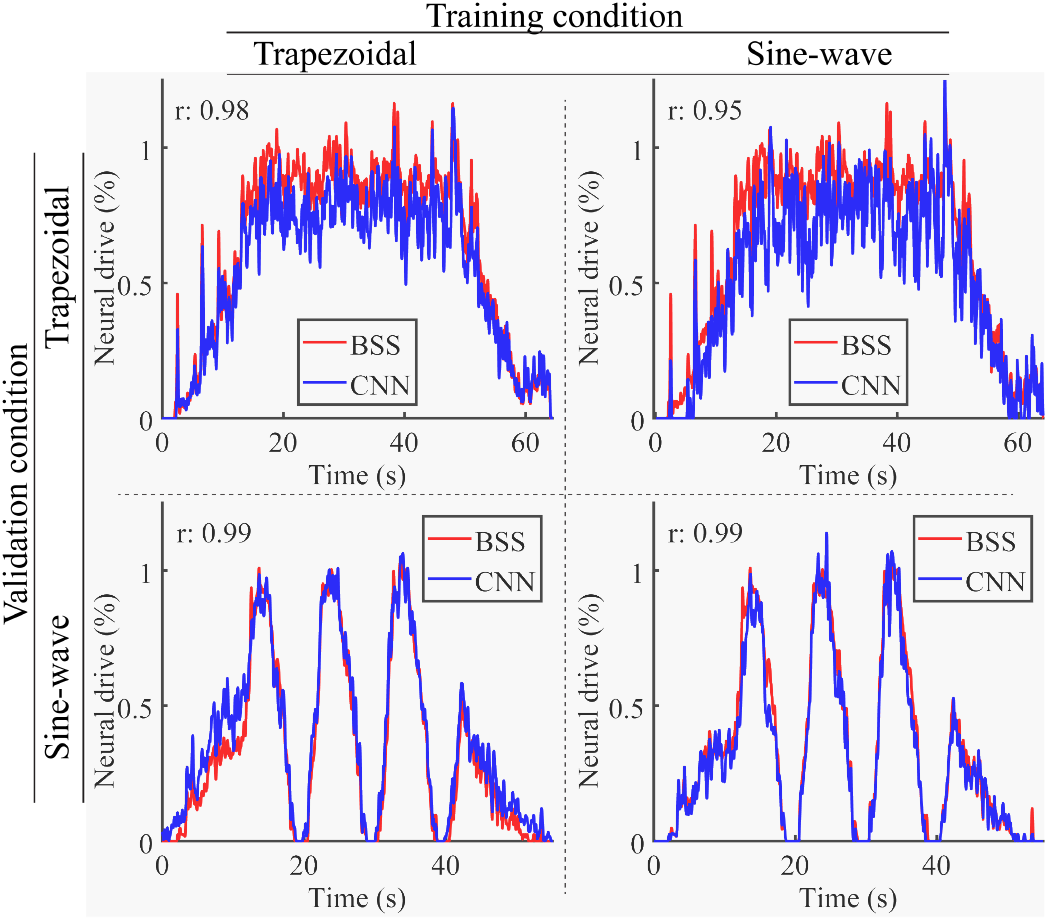
A comparison example between neural drive from the blind source separation (BSS) and neural drive from the deep CNN trained with different datasets. For column 1 (a and d), the deep CNN was trained with data from 20% MVC; for column 2, the deep CNN was trained with data from sinusoidal torque tracking; for column 3, the deep CNN was trained with combined data from 20% MVC and sinusoidal torque tracking.

**Fig. 6:**
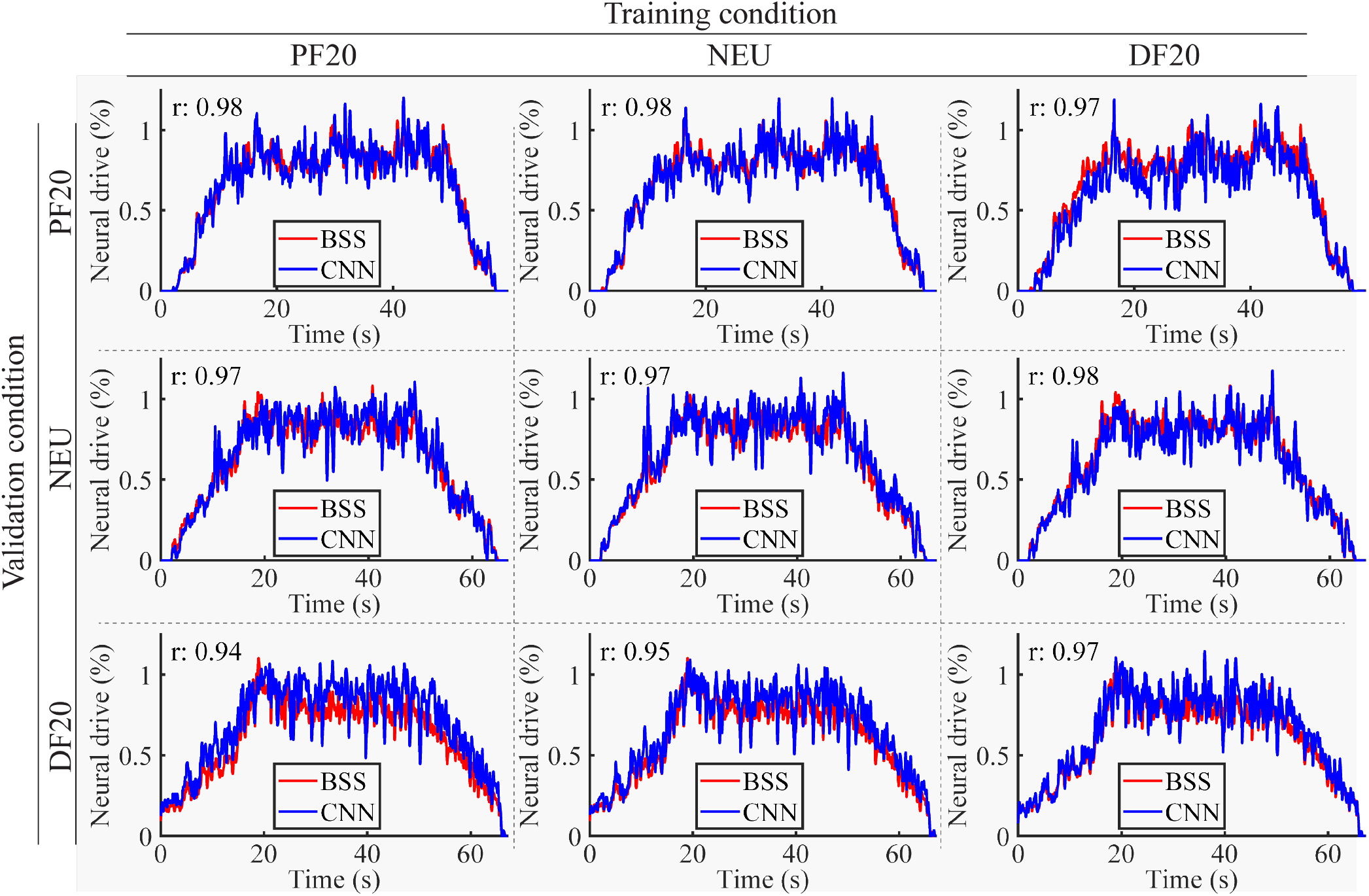
A comparison example between neural drives estimated with the blind source separation (BSS) and with the deep CNN trained with different datasets. For column 1, the deep CNN was trained with data from PF20; for column 2, the deep CNN was trained with data from NEU; for column 3, the deep CNN was trained with data from DF20. From row 1 to 3, the data for the validation were recorded during PF20, NEU, and DF20 contraction tasks.

## IV. Discussion

We aimed at developing a novel deep CNN framework for neural drive estimation in the form of CST from HD-EMG signals. The proposed deep CNN was trained and validated with HD-EMG signals and the corresponding CST extracted using the BSS method (gold standard) from three scenarios: 1) isometric trapezoidal contractions with different intensities, 2) isometric contractions tracking trapezoidal and sinusoidal torque targets, 3) isometric trapezoidal contractions with different ankle angles. We found that the input and output structures of the deep CNN have significant effects on the performance of the neural drive estimation (Table. II), which aligns with previous studies [26]. This is also supported by the fact that the window size affects the number of MUAPs included in the HD-EMG signals and corresponding MU firing activities, which is related to the number of outputs. The main results (Table. III, IV, and V) suggest that the proposed deep CNN framework is: 1) capable of extracting spatio-temporal features from HD-EMG signals and accurately estimating the neural drive (intra-task validation), 2) generalizable across different contraction tasks within each scenario (inter-task validation), and 3) scalable to complex datasets including multiple contraction tasks (last column in Table. III, IV, and V).

The deep CNN generated slightly lower correlation coefficients and higher nRMSE in inter-task validations than in intra-task validations, and such performance decrease might be affected by the training data. We speculate that such performance inconsistency might be explained by the difference in the number of MUs and in the MUAP shapes between the training and the testing data. Specifically, when performing repeated contractions for the same contraction tasks within a short period of time, same MUs decomposed using BSS resulted in similar MUAP shapes. Thus, the deep CNN generated higher correlation coefficient and lower nRMSE in intra-task validations. However, for inter-task validations, the number of identified MUs and their MUAP shapes were different across different contraction tasks (Table. I). First, the decomposition algorithm converged towards the MUs that contribute more to the signal, i.e., the biggest MUs [32]. Thus, these MUs changed across intensities. Second, the ankle angle change caused changes in muscle length and pennation angle as well as the tissues between the muscle and the electrode [33]. These changes contributed to relative shift between muscle fibers and electrodes and consequently caused changes in the MUAPs shapes (e.g., amplitude and duration) and their distribution across the measurement channels. In summary, the discrepancy of the MUAP shapes included in the training data and testing data deteriorated the performance of the deep CNN slightly in the inter-task validations.

We noticed that the neural drive estimated from deep CNN was jerkier than that from human-edited spike trains (Fig. 5 and Fig. 6). One potential reason is that the human operator manually edited the spike trains to add false negatives and remove false positives [29]. The high frequency components in the estimated neural drive, however, might not be an issue for human-robot interfacing, since a low-pass filter is typically used in EMG-based controllers to remove undesired high frequency control signals so as to generate smooth and stable movement commands for wearable robots [34], [35]. Therefore, with proper filtering, the deep CNN framework is a promising approach to achieve real-time human-machine interfacing for wearable robots. We will investigate these factors in our future studies.

In this study, one limitation is that we only demonstrated the feasibility of the deep CNN under three representative scenarios with specific contraction intensities, force profiles, ankle angles and with limited participants. Although we speculate the findings will apply to different levels of contraction intensities and muscle length, our future studies will investigate the feasibility of the proposed deep CNN under dynamic contractions with varying levels of muscle activation and muscle length. In parallel, another future work is to investigate the feasibility of the deep CNN using training data based on simulated HD-EMG signals for dynamic contractions and different subjects. This would allow us to train and validate the CNN with various contraction tasks (i.e., activation intensities and muscle lengths) with minimal labor-intensive manual edits and experimentation. In addition, we will attempt to apply the trained deep CNNs with simulation data to experimental data. The success of this approach could potentially create a general deep CNN that can handle other possible situations for reallife applications.

Human-robot interfacing relies on two steps to generate control commands. The first step consists of extracting the neural drive to each muscle from HD-EMG, and the second step consists of combining the neural drive information from these muscles to generate the control command at the joint level. Another limitation of this study is that we only addressed the first step to estimate the neural drive from HD-EMG under different contraction tasks using a unified deep CNN. In our future work, we will combine neural drives from multiple muscles with their mechanical properties or force output to generate control commands (e.g., torque) for human-robot interfacing.

## V. Conclusion

We proposed a deep CNN framework to estimate the neural drive in the form of CST from HD-EMG signals across different isometric contraction tasks. Our study demonstrated that 1) the deep CNN is generalizable to different contraction tasks within each scenario, i.e., estimating the neural drive for contraction tasks that are not included in the training data, and 2) the deep CNN is scalable to capture complex features of variant MUAP shapes when trained with combined data from different contraction tasks, which also increases the accuracy of the neural drive estimation (last column in Table. III, IV, and V). Comparing with the commonly used offline neural drive extraction approach (i.e., BSS), the proposed deep CNN framework could generate accurate estimation of the neural drive. Moreover, the proposed work is generalizable to different contraction tasks without retraining and could run in a realtime manner as was done in a previous study [26]. Therefore, the proposed deep CNN framework is a promising candidate for real-time human-machine interface based on neural drive for assistive technology (e.g., exoskeletons and prostheses).

